# Multiscale Network Disruption of Auditory-Language Circuits Underlying Auditory Hallucinations in Schizophrenia

**DOI:** 10.64898/2026.10.05.756754

**Authors:** Andrew C Murphy, Antigona Martinez, Yadi Chen, Michael Avissar, Bahar Hazal Yalçınkaya, Juan Sanchez-Pena, Gaurav H Patel, Daniel C Javitt

## Abstract

**Background:** Auditory verbal hallucinations (AVH) are among the most disabling symptoms of schizophrenia, yet the network-level mechanisms linking auditory dysfunction to hallucination severity remain poorly characterized, limiting targeted therapeutic development.

**Methods:** Using resting-state fMRI and high-resolution surface-based parcellation (HCP-MMP), we examined auditory-language functional connectivity and mesoscale network organization in 56 patients with schizophrenia and 22 healthy controls (primary dataset, N=78). Pairwise auditory-language connectivity associations with hallucination severity (PANSS P3) were examined alongside auditory network community structure, assortativity, hub organization, and routing efficiency. To further characterize these parcel-level findings at the vertex level, we applied a region-growing approach to identify auditory-frontal connectivity patterns maximally associated with severity. In silico perturbation analyses tested the causal role of belt cortex connectivity. Key findings were independently replicated in 93 patients from the Human Connectome Project for Early Psychosis (HCP-EP), for a combined total of 171 participants.

**Results:** Auditory-language connectivity disruptions were concentrated in belt cortex and specifically implicated ventral what-stream connections, with a vertex-wise analysis converging on the same anatomy. Auditory network segregation scaled with hallucination severity, with belt cortex emerging as the primary integrative hub governing auditory-to-language routing efficiency. Perturbation analyses showed that strengthening earlier-hierarchy auditory regions reduced network segregation more than strengthening later-hierarchy regions. Reduced belt cortex coupling and elevated network segregation replicated independently in the HCP-EP sample.

**Conclusions:** Auditory network disorganization scales with hallucination severity and replicates across independent samples, with belt cortex as the critical locus of connectivity disruption and network integration failure, identifying it as a candidate neuromodulatory target.

## Introduction

Auditory verbal hallucinations (AVH), hearing voices in the absence of external speech, are among the most prevalent and disabling symptoms of schizophrenia, affecting most patients and persisting despite antipsychotic treatment in up to a third of cases [1]. Two regions are central to this process: auditory cortex, which processes sound from basic spectral features to complex auditory objects, and language cortex, which processes, monitors, and produces speech. Coordination between them is thought critical for distinguishing internally generated from externally perceived speech, and its breakdown may underlie AVH [2]. Resting-state fMRI functional connectivity (FC) studies of hallucination severity have consistently implicated reduced auditory-language coupling [3, 4], converging with structural findings that AVH severity correlates most strongly with left superior temporal gyrus and insula atrophy [5]. However, connectomic-level analysis of AVH mechanisms has been limited by the coarseness of traditional atlases: the Brodmann map divides auditory and posterior language cortex into just three regions (primary auditory cortex, area 41; secondary auditory cortex, area 42; Wernicke’s area, 22), with anterior language cortex divided only into Broca’s areas 44 and 45.

High-resolution surface-based parcellation now addresses this gap: the Human Connectome Project multimodal parcellation atlas (HCP-MMP; [6]) divides auditory cortex into 8 parcels based on myelin content, architecture, task-based fMRI response, and functional connectivity. We assigned these parcels to functional networks using the Cole–Anticevic network partition [7], which also defines a 14-parcel language network across posterior and anterior regions (Fig. 1E). Effective-connectivity studies reveal a hierarchical organization from primary auditory cortex (A1) through midlevel belt regions (LBelt, MBelt, PBelt) to higher-level secondary auditory cortex (A4), from which two distinct processing streams emerge [8, 9].

**Figure 1:**
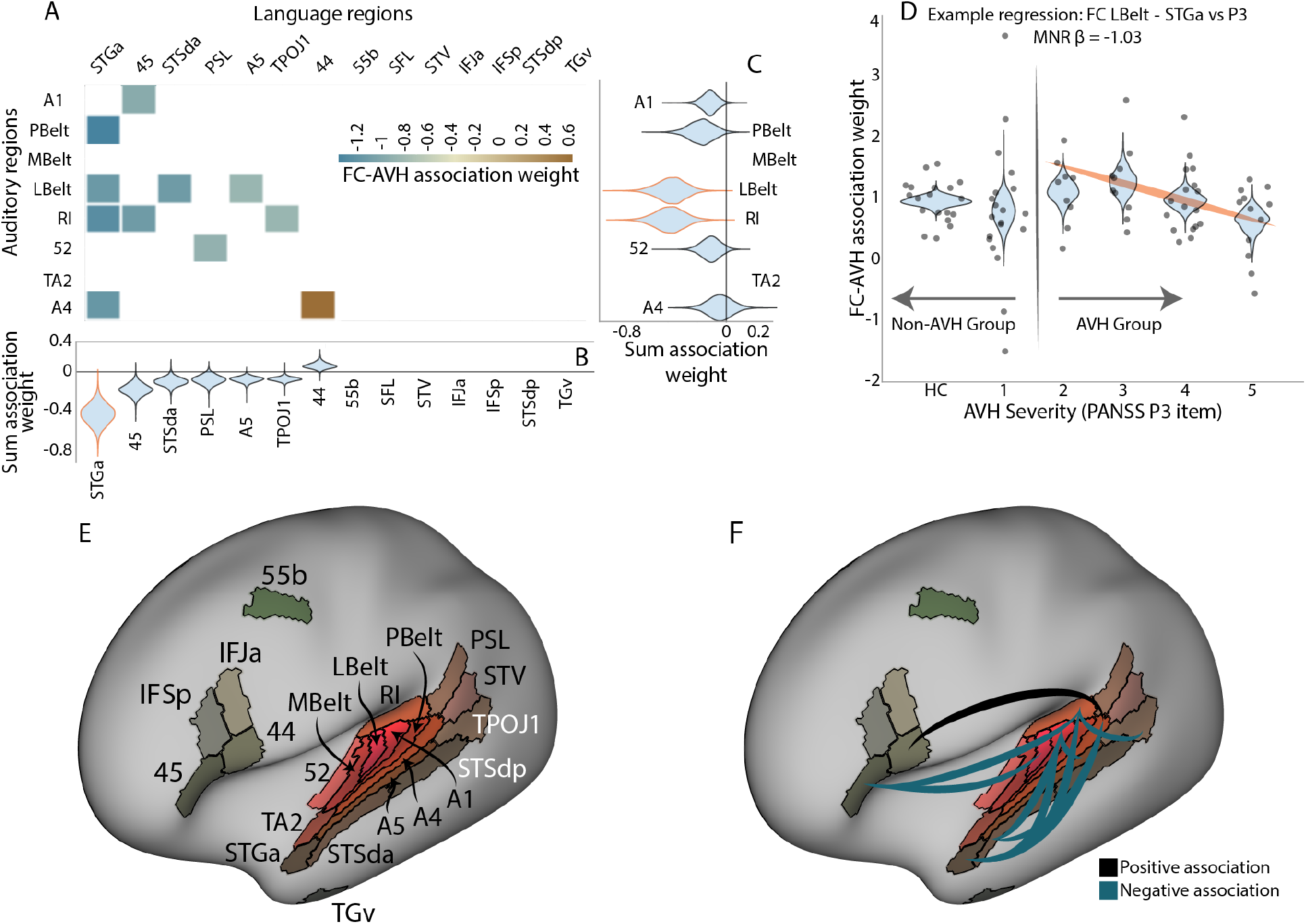
Parcel-to-parcel auditory–language functional connectivity analyses reveal belt-concentrated severity-linked disruptions. (A) Multinomial regression coefficients (*β*) for auditory–language FC associations with AVH severity (PANSS P3); negative values indicate lower FC at higher severity. (B) Aggregate incident weights for language parcels, showing STGa as the dominant language target; violins show boot-strapped group-wise mean distributions. (C) Same as (B) for auditory parcels, showing RI and LBelt as the dominant auditory contributors. (D) Example multinomial regression for the LBelt–STGa connection: columns correspond to HC and PANSS P3 scores 1–5, with a divider between columns 2 and 3 demarcating non-hallucinators from hallucinators; individual FC values are shown as swarm plots over bootstrapped violin plots, with a line of negative slope connecting hallucinator columns 3–6. (E) Labeled cortical surface showing the auditory and language parcels included in the analysis. (F) Cortical surface rendering of severity-linked auditory–language FC disruptions; belt auditory cortex, and LBelt in particular, shows the greatest concentration of severity-linked FC reductions with language regions. Line thickness indicates regression coefficient magnitude; blue connections indicate negative associations (lower FC with greater severity), black indicate positive associations. All associations are FDR-corrected and spin-test adjusted.

From A4, a ventral “what” stream progresses through temporal sulcus regions to language cortex, supporting auditory object identification: who is speaking and what is said, while a dorsal “where” stream projects to superior parietal regions for spatial processing and sensorimotor integration. Disruptions to this hierarchy have been associated with AVH in schizophrenia [9]. Complementing this hierarchical view, network-level perspectives consider pairwise couplings jointly across all areas, forming a densely connected system whose emergent architectural properties may also be disrupted in schizophrenia.

We have previously used network-level connectomic measures including community detection and assortativity, which characterize how sharply networks organize into densely interconnected modules [10], and participation coefficient, which identifies hub regions supporting centralized routing and efficient long-range communication [11–13], to define subnetworks underlying working memory in healthy volunteers [14]. Here we apply these tools to auditory connectivity disturbances relative to AVH severity, using pairwise and mesoscale approaches across two independent datasets totaling 171 participants. We hypothesized that reduced early auditory-language connectivity would relate to altered community structure and routing efficiency, which would in turn correlate with AVH severity, and tested this using virtual lesion and repair perturbation analyses. Because neuromodulatory approaches to AVH have been only partially effective, in part due to coarse anatomical targeting, we use parcel-level connectivity analysis to refine existing targets.

## Methods

### MRI Processing

MRI acquisition and preprocessing are described in the Supplementary Methods. Briefly, structural and funcstional MRI data were preprocessed using fMRIPrep [15, 16] and post-processed using XCP-D [17]; functional brain networks were constructed from Fisher *r*-to-*z* transformed Pearson correlation coefficients between parcel time series from the Glasser multimodal cortical atlas [6].

### Statistical analyses

Associations between network metrics and AVH severity were assessed using multinomial regression with PANSS P3 as the outcome, with significance evaluated using nonparametric permutation testing (10,000 permutations) and FDR-corrected multiple comparisons. Spatial autocorrelation was accounted for using spin-test null models [18]; full statistical details are in the Supplementary Methods.

### Network metrics

Auditory network community structure was identified for each subject using modularity maximization (Louvain algorithm), partitioning the network into groups of densely interconnected regions. From this partition, we computed *assortativity*, measuring how sharply segregated the communities are, with higher values indicating greater internal fragmentation. We computed *pathway efficiency* using mean first passage time from random-walk dynamics, capturing how efficiently signals travel from auditory source regions through belt cortex to language targets. For each parcel, we computed an *incident total-weight* measure summarizing the cumulative strength of its significant connections, and a *participation coefficient* quantifying how much a region’s connections span multiple communities, distinguishing hub regions bridging communities from those confined to one community. Full mathematical definitions are in the Supplementary Methods.

### Incident total-weight measures

The incident total weight for each parcel was defined as the sum of all significant (FDR- and spin-test-corrected) multinomial regression coefficients for edges incident on that parcel. Pairwise differences between parcel incident weights were assessed using permutation testing in which parcel labels were randomly shuffled to generate null distributions.

### Participation coefficient

The participation coefficient [19] of parcel *i* was defined as:

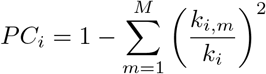

where *k*_*i*_ is the total weighted degree of parcel *i* and *k*_*i,m*_ is the sum of edge weights to parcels in community *m*. Values approaching 0 indicate provincial hubs; values approaching 1 indicate connector hubs. Associations between participation coefficients and auditory network assortativity were assessed using Pearson correlation with permutation-based significance testing, and pairwise differences between parcels were evaluated using permutation-based pairwise tests.

### Vertex-wise region-growing analysis

To identify auditory–frontal functional connectivity patterns maximally correlated with hallucination severity, we applied a vertex-wise region-growing algorithm within predefined left-hemisphere search spaces spanning auditory cortex and inferior frontal cortex (HCP-MMP1.0 parcellation). For each participant with PANSS P3*>*1, we computed pairwise FC between all auditory and inferior frontal vertices and correlated these with P3 across participants. A greedy region-growing algorithm then identified spatially contiguous vertex clusters, one anchored in auditory cortex, one in inferior frontal cortex, whose mean interconnecting FC showed the strongest positive and negative correlation with AVH severity. Clusters were required to reach a minimum surface area of 200 mm^2^, chosen to ensure sufficient spatial extent for effective HD-tDCS current delivery. This procedure was applied separately for positive and negative FC–severity correlations, yielding two pairs of auditory–frontal regions whose connectivity most strongly tracks hallucination severity in opposite directions.

### Ethical approval

All study procedures were approved by the Institutional Review Board of the Nathan Kline Institute for Psychiatric Research. All participants provided written informed consent prior to enrollment, and all experiments were conducted in accordance with the Declaration of Helsinki. Replication analyses used data from the Human Connectome Project for Early Psychosis (HCP-EP), details are provided elsewhere [20].

## Results

### Participants

The primary dataset included 22 healthy controls (HC; mean age 36.6 *±* 10.2 years, 55.6% male) and 56 individuals with schizophrenia (mean age 37.5 *±* 10.5 years, 79.7% male; 16 non-hallucinating, PANSS P3=1; 40 hallucinating, PANSS P3*>*1). The replication dataset, HCP-EP, comprised 93 individuals with schizophrenia spectrum disorder (mean age 22.6 *±* 3.6 years, 59% male; 66 non-hallucinating, 27 hallucinating). Inclusion and exclusion criteria and acquisition parameters are in the Supplementary Methods.

### Parcel-to-parcel rsFC

Building on prior reports of reduced auditory-language coupling in AVH, we first examined this relationship at the level of individual parcel pairs. In an initial left auditory-to-left language parcel analysis, 11 connections survived FDR and spin-test correction (Fig. 1A; example in Fig. 1D), shown on a labeled cortical surface in Fig. 1E. To assess specificity, we repeated the analysis for left auditory-to-left somatomotor connections, with no connections surviving, and for right auditory-to-right language connections, where a single connection survived (right TA2 to right IFJa, *β* = 0.90, *p <* 0.001). These associations were specific to AVH severity: repeating the analysis using PANSS positive (excluding P3), negative, and general psychopathology subscales yielded no significant connections. To quantify subfield specificity, we computed incident total-weight measures for each parcel (Fig. 1B,C), shown on the cortical surface in Fig. 1F. RI and LBelt showed the largest-magnitude negative incident weights of all auditory parcels. A vertex-wise region-growing analysis confirmed this pattern: the auditory–frontal connection most negatively correlated with AVH severity linked a cluster spanning posterior belt cortex (A1, LBelt, MBelt, area 52) to inferior IFSp and superior area 45 (Fig. 2C, blue).

**Figure 2:**
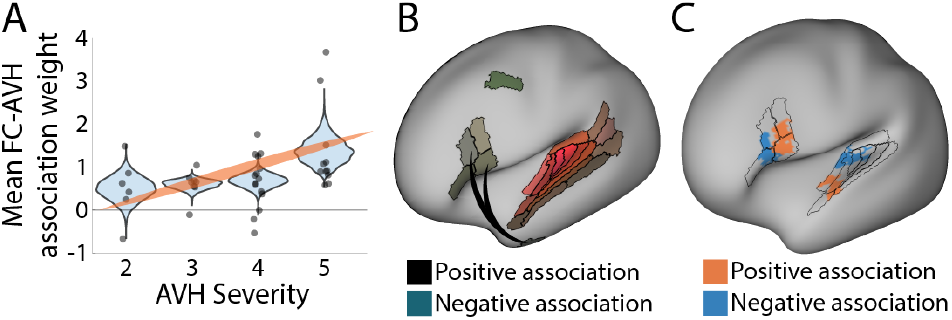
Language-to-language functional connectivity (FC) disruptions and their relationship to A4 connectivity. (A) Mean functional connectivity of the two significant language-to-language connections shown in (B) increases with AVH severity (multinomial regression, *β* = 1.2202, *p <* 0.001); points are individual values, violins show bootstrapped group-wise mean distributions for PANSS P3 scores 2–5. (B) Language-to-language FC associations with AVH severity. Parcel colors correspond to the Cole–Anticevic language network parcellation [7]. (C) Vertex-wise region-growing analysis identifying auditory–frontal connectivity patterns maximally tracking AVH severity; blue indicates the connection most negatively correlated with severity, linking posterior belt cortex (A1, LBelt, MBelt, area 52) to inferior IFSp and superior area 45; orange indicates the connection most positively correlated with severity, linking anterior auditory cortex (area 52, MBelt, A4, posterior TA2) to inferior IFJa and superior area 44.

A4 showed a mixed pattern at the parcel level, with increased FC to area 44 but decreased FC to STGa at higher AVH severity. Within the language network, STGa showed the largest-magnitude negative incident weight, uniquely exceeding all other parcels. Aggregate FC between the top auditory parcels (LBelt, RI) and STGa was significantly lower in high-relative to low-severity hallucinators (Δ = 0.54, *p* = 0.027), and significantly closer to low-than high-severity hallucinators in healthy controls (ΔΔ = 0.54, *p* = 0.028). The vertex-level region-growing analysis confirmed this positive association: the auditory–frontal connection most positively correlated with AVH severity linked anterior auditory cortex (area 52, MBelt, A4, posterior TA2) to inferior IFJa and superior area 44 (Fig. 2C, orange).

Extending this analysis to language-to-language connections between anterior and posterior regions, two connections survived FDR and spin-test correction (Fig. 2B): area 44–TGv and IFSp–TGv, both showing increased FC with worsening AVH; mean FC across these two connections scaled significantly with severity (multinomial regression, *β* = 1.2202, *p <* 0.001; Fig. 2A).

### Mesoscale analyses

Having found that auditory-language decoupling tracks AVH severity, we next asked whether one side of the circuit—auditory or language—is more responsible for this reorganization. We used a community-detection approach similar to our prior study [14] to partition each network into communities and compute assortativity. Auditory network assortativity scaled with AVH severity (multinomial regression, *β* = 1.64, *p* = 0.041; Fig. 3A), with more segregated communities relating to more severe hallucinations, whereas language network assortativity was unrelated to severity (*β* = 0.26, *p* = 0.417), indicating that this architectural reorganization is specific to the auditory system. This association was specific to hallucination severity, as it was unrelated to other positive symptoms (PANSS positive excluding P3), negative symptoms, or general psychopathology. Auditory assortativity was also negatively correlated with overall auditory-to-language FC (*r* = *−* 0.47, *p* = 0.006), linking greater community segregation to greater auditory-language decoupling.

**Figure 3:**
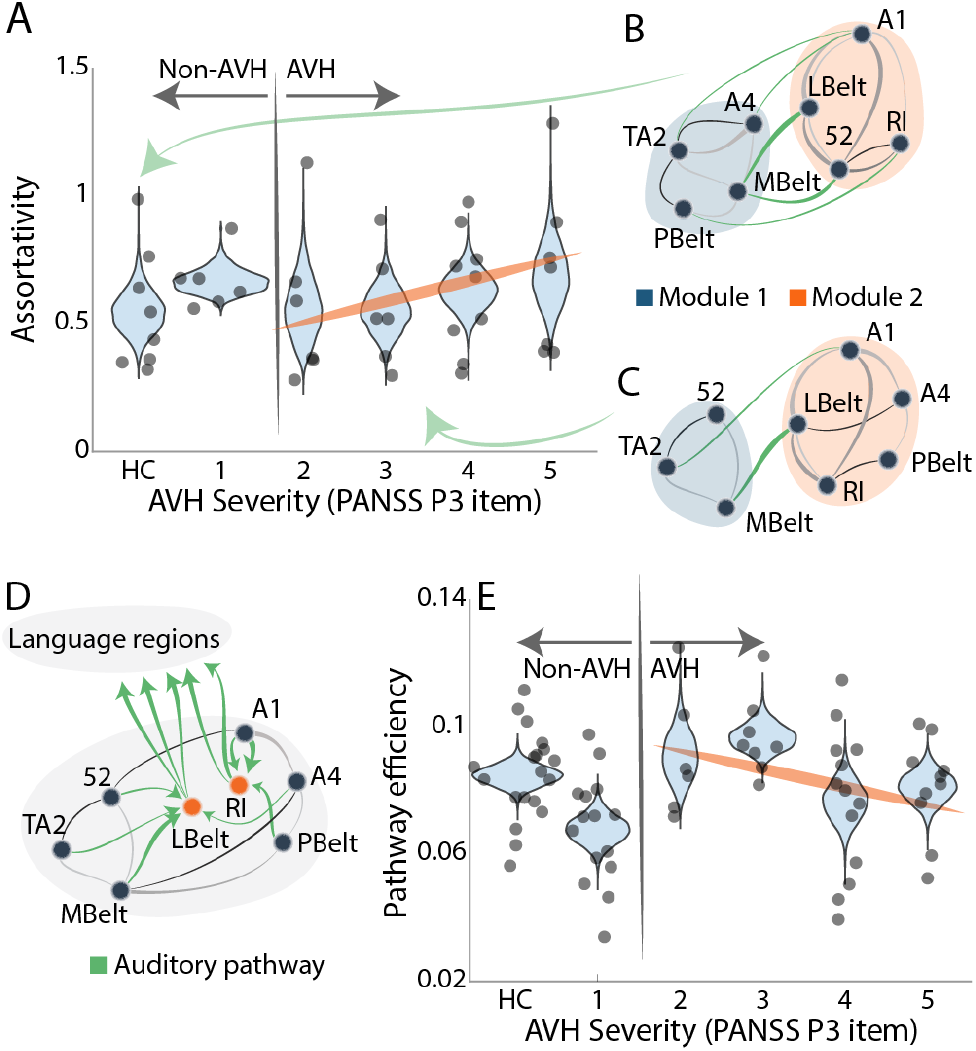
Auditory network segregation and routing inefficiency scale with auditory verbal hallucination (AVH) severity. (A) Auditory network assortativity increases with AVH severity (multinomial regression, *β* = 1.64, *p* = 0.041); points are individual values, violins show bootstrapped group-wise mean distributions. Example of an integrated auditory network architecture observed in healthy controls. (C) Example of a segregated auditory network architecture observed in schizophrenia patients. (D) Schematic of the auditory– language communication pathway from auditory source parcels through LBelt/RI to language target region STGa. (E) Reduced auditory–language pathway efficiency predicts greater AVH severity (multinomial regression, *β* = *−*20.29, *p* = 0.021).

To understand this auditory-specific reorganization in more detail, we examined its underlying community structure, hub organization, and routing efficiency. In healthy controls, the auditory network divided into a superior, largely early hierarchical community (A1, LBelt, 52, RI) and an inferior, predominantly late hierarchical community (A4, MBelt, PBelt, TA2) (Fig. 3B); the weighted stochastic block model (WSBM; *k* = 2) [14,21,22] yielded identical consensus community assignments in both groups, supporting the stability of this two-community solution. In hallucinating participants, these communities reorganized: area 52 moved from the early into the later, higher-association community, while A4 and PBelt moved oppositely (Fig. 3C).

Modular separation is influenced in part by integrative hubs. Quantifying each auditory parcel’s participation coefficient, LBelt emerged as the dominant hub, with its integrative capacity more strongly associated with auditory network integration than any other parcel (*p* = 0.016); the analogous test for RI was not significant (*p* = 0.889), despite RI’s comparable contribution to auditory-language FC disruption.

Reduced efficiency of the auditory *→* LBelt/RI *→* STGa pathway (schematic in Fig. 3D) was associated with higher AVH severity (multinomial regression, *β* = *−*20.29, *p* = 0.021; Fig. 3E), exceeding a randomized-pathway null (*p* = 0.007). This pathway’s directionality reflects the established feedforward hierarchy of auditory processing [8, 9]; LBelt, RI, and STGa were selected given their dominant incident weights in the parcel-level analysis (Fig. 1B,C). This association was likewise specific to hallucination severity (all other PANSS domains: *p >* 0.05). Pathway efficiency showed a trend-level negative correlation with auditory network assortativity (*r* = *−* 0.29, *p* = 0.064) that did not exceed a randomized-pathway null (*p* = 0.777), suggesting these measures capture partially independent aspects of auditory network organization.

### Perturbation analyses

Finally, we sought a complementary perspective via an *in silico* perturbative model: for each auditory parcel, we constructed a continuous spectrum of connection scaling from weakening (HC cohort) to strengthening (AVH+ cohort), examining the relationship between this spectrum and resulting changes in auditory network assortativity and pathway efficiency. Strengthening earlier-hierarchy nodes (e.g., LBelt, RI) integrated the auditory network significantly more than strengthening later-hierarchy nodes (Spearman *r* = 0.73, *p* = 0.023). Focusing on LBelt and RI, this spectrum tracked monotonically with pathway efficiency: efficiency declined as connectivity weakened and increased as it strengthened.

### Replication in an independent sample

To assess replicability, key findings were tested in an independent sample from the HCP-EP dataset. The LBelt auditory-language incident weight was significantly lower in the AVH+ group than the AVH-group, relative to the distribution of group differences across all auditory-language connections in the network (*diff* = *−* 0.051, one-tailed rank *p* = 0.044; Fig. 4A), replicating the primary finding. Auditory network assortativity, computed using the primary dataset’s consensus community partition, was significantly higher in hallucinating participants (diff = 0.27, *p <* 0.001; Fig. 4B), replicating the finding of greater auditory network segregation in AVH. Auditory-to-language pathway efficiency was lower in hallucinating participants in the expected direction but did not reach significance.

**Figure 4:**
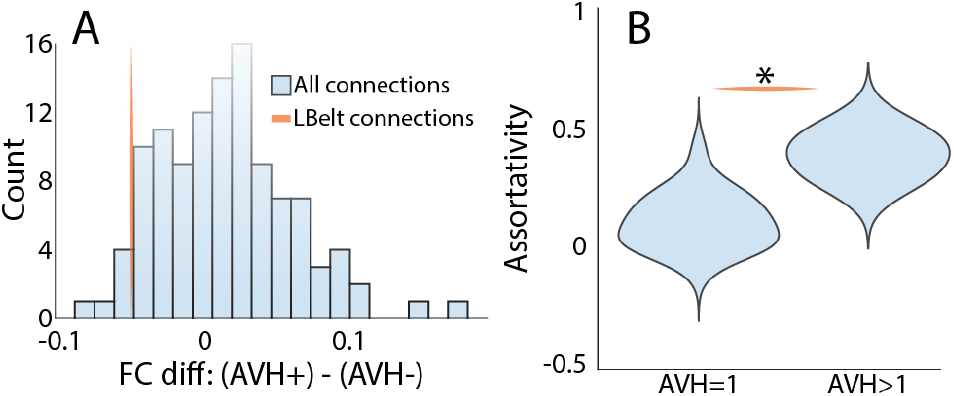
Replication of key findings in an independent Human Connectome Project - Early Psychosis (HCP-EP) sample. (A) The LBelt auditory-language incident weight difference between AVH+ and AVH-participants (diff = *−* 0.051 ) is significantly larger in magnitude than expected relative to the distribution of group differences across all auditory-language connections (one-tailed rank *p* = 0.045); bars show the distribution of group differences across all auditory-language connections, line indicates the observed difference for the LBelt connection. (B) Auditory network assortativity, computed using the consensus community partition derived from the primary dataset, is significantly higher in hallucinating participants (diff = 0.27, *p <* 0.001).

## Discussion

Brain network architecture - how regions form communities, how hubs route information between them, and how efficiently signals transmit - determines network capacity for information processing, offering insights pairwise connectivity alone cannot capture. Here, we apply this framework to the auditory system in schizophrenia, evaluating both parcel-wise and mesoscale structure relative to AVH severity.

### Parcel-level FC reveals hierarchical specificity of auditory-language coupling in AVH

Prior studies of auditory-network coupling in AVH have largely relied on seed-based volumetric approaches, characterizing disruptions at the level of broad anatomic regions such as Broca’s or Wernicke’s areas [3, 4]. In contrast, our parcel-wise auditory-language network analysis localized disruptions to belt cortex and ventral “what”-stream connections specifically, suggesting relative “what”-stream underrepresentation in severe AVH. Belt cortex occupies the hierarchical interface between auditory and language systems [8], a natural point for auditory-language communication to concentrate and fail; predictive coding accounts suggest that disruption here leaves top-down predictions poorly constrained by bottom-up input [23–25], while auditory-deafferentation accounts predict a similar breakdown [26, 27]. This is consistent with prior findings linking early auditory processing measures such as mismatch negativity to hallucination severity [28–33], and with the dual-stream account in which selective “what”-stream disruption impairs auditory object identification and voice source attribution [8, 34]. Critically, a vertex-wise region-growing analysis restricted to auditory-to-inferior-frontal connections, but otherwise agnostic to parcel boundaries, converged on the same anatomy, linking posterior belt cortex to ventral inferior frontal cortex as the connection most negatively tracking AVH severity. The identification of IFSp and area 45 as the frontal endpoint is consistent with Rolls and colleagues’ characterization of these regions as key nodes within the human language effective connectome, receiving auditory what-stream input from A5 and STGa [35]. This convergence across methodologically independent approaches strengthens confidence that the disruption reflects genuine functional organization rather than parcellation-dependent artifacts; the parcel-based framework is further motivated by enabling computation of mesoscale network metrics not accessible at the vertex level.

A notable exception is A4, which showed increased FC with where-stream inferior frontal region area 44 at higher AVH severity at the parcel level. A4 is the hierarchical bifurcation point beyond which the dorsal and ventral auditory streams diverge [8], and this positive association suggests where-stream overrepresentation compounding the what-stream underrepresentation described above; a relative shift in auditory processing toward spatial and sensorimotor integration at the expense of object identification. The vertex-wise region-growing analysis again converged on this pattern, identifying a broader anterior auditory cluster spanning area 52, MBelt, A4, and posterior TA2 as the region whose frontal connectivity most positively tracked AVH severity, with the frontal endpoint spanning inferior IFJa and superior area 44. That both the parcel-level and vertex-level approaches independently implicate anterior auditory cortex and dorsal-stream frontal regions in the positive association, without the vertex-wise analysis being constrained to A4 or area 44 specifically, strengthens the interpretation that where-stream connectivity is genuinely overrepresented relative to what-stream connectivity in severe AVH. Specificity was further supported by right hemisphere analyses, where only a single connection survived: right TA2 to right IFJa, paralleling the left-lateralized what-stream pattern though far more limited in extent.

The two significant language-to-language associations, area 44–TGv and IFSp–TGv (both increasing with worsening AVH) show a similar pattern: TGv is itself a ventral “what”-stream region [8], yet its connectivity increased with both a “where”-stream (area 44) and “what”-stream (IFSp) partner as symptoms worsened, paralleling the bifurcation-point disruption at A4 and suggesting a similar loss of stream segregation down-stream in language cortex.

### Network architecture reveals belt cortex as cross-module linking hub and routing bottleneck

Moving beyond pairwise connections, we next examined how AVH severity relates to emergent mesoscale properties of the auditory network: community structure, hub organization, and routing efficiency. These properties relate to behavioral outcomes in healthy populations [14,36,37] and are disrupted in schizophrenia [38–40]. In HC, the auditory network organized into two communities: a core community of MGN-recipient regions (A1, LBelt, RI, area 52) and a higher-association community (TA2, MBelt, A4, PBelt), broadly consistent with the hierarchy described by Rolls and colleagues [8, 9]. In the AVH cohort, these communities reorganized: area 52 moved into the higher-association community, while A4 and PBelt moved oppositely, blurring the boundary between earlier and later auditory processing at the same point where the two streams diverge.

At the whole-brain level, both modularity [40] and community structure [41] may be disrupted in schizophrenia. In the auditory system specifically, greater assortativity (sharper internal fragmentation) was associated with more severe hallucinations, consistent with disconnection accounts of schizophrenia [42, 43]. Excessive segregation may limit information export beyond what connection strengths alone predict [44–46]. Auditory assortativity was also negatively correlated with auditory-language FC, suggesting the coupling deficits described above partly reflect architectural constraints at the community level.

This community-level segregation is in part determined by cross-community linking hubs [47,48], which at the whole-brain level are abnormal in schizophrenia [44]. LBelt emerged as the dominant integrative node of the auditory network, its participation coefficient more strongly associated with auditory network integration than any other region, consistent with structural connectivity studies showing LBelt has particularly broad connectivity within the auditory system compared to other belt regions [9,49]. Notably, this finding is specific to LBelt rather than RI, which contributes substantially to auditory-language FC disruption but does not show the same dominant relationship with community segregation.

Reduced hub-mediated integration may also impair routing efficiency, since hub organization, particularly rich-club connectivity among hubs, provides the structural basis for long-range routing [44, 48]. Reduced auditory-to-language routing efficiency through belt cortex was specifically associated with hallucination severity: as belt cortex loses its integrative function, signals become less available downstream for the top-down predictions and source-monitoring that normally prevent internally generated activity from being experienced as external voices [23, 27, 50]. Pathway efficiency and assortativity were only weakly related, indicating partially independent organizational features. Together, community structure, hub integrity, and routing efficiency offer three complementary views of the same disruption.

### Perturbation analyses and clinical implications for neuromodulatory intervention

These correlational findings across community structure, hub integrity, and routing efficiency all implicate belt regions in auditory network dysfunction. To move beyond correlation, we conducted *in silico* perturbation analyses following the connector hub lesioning framework [47, 51]: strengthening earlier-hierarchy nodes integrated the auditory network significantly more than strengthening later-hierarchy nodes, indicating early auditory regions exert outsized control over network organization. Strengthening LBelt connectivity specifically also increased pathway efficiency, suggesting integration and routing efficiency are mechanistically linked.

Non-invasive brain stimulation, including TMS and tDCS, has shown promise for persistent AVH, though results have been variable [52], in part due to coarse anatomical targeting. LBelt emerges as a primary candidate: it is the dominant integrative hub, shows the strongest auditory-language FC disruption, and perturbation analyses suggest strengthening early-hierarchy connectivity normalizes network organization, converging with prior evidence implicating LBelt via intrinsic neural timescales [9] and MGN input disruption [28]. A temporoparietal coordinate used in several imaging-guided rTMS studies [53] corresponds anatomically to RI [54, 55], suggesting these studies may have targeted belt cortex; surface-based localization of LBelt and RI offers a principled basis for refining this target, since variable localization has been proposed as a key source of mixed efficacy [55]. The anterior auditory cluster identified by the vertex-wise region-growing analysis represents a second, accessible candidate for TMS or high-definition tDCS.

### Replication in an independent sample

The primary findings were replicated in an independent HCP-EP sample. As in our primary analysis, LBelt auditory-language coupling was reduced and auditory network assortativity was elevated in hallucinating participants. The assortativity finding was particularly prominent in the early-psychosis replication dataset, which contained younger participants than our primary data set. Future studies extending the analyses to even earlier samples, such as prodromal populations, might therefore be warranted. That these effects replicate across datasets, using community structure derived from the primary cohort, supports the generalizability of both the parcel-level FC disruption and mesoscale organizational findings. Pathway efficiency trended in the expected direction but did not reach significance, consistent with reduced power in a binary group comparison relative to the continuous severity analysis in the primary dataset.

## Conclusions

In humans, auditory processing ascends hierarchically from primary auditory cortex through belt regions and into parallel “what” and “where” pathways that process content and location of speech independently. Here, using parcellations derived from the HCP-MMP atlas, we show that the normal modularity of early auditory processing is disrupted at the level of the auditory belt regions, leading to a reduction in assortativity within the early auditory system in hallucinating vs. non-hallucinating individuals, and disruption of “what” vs. “where” modularity. Both reduced correlation between belt and posterior language regions, and increased correlation between posterior and anterior language regions correlated with greater hallucisnation severity. These findings converge with other evidence of overreliance on top-down vs. bottom-up information in schizophrenia. In perturbation analyses, strengthening connectivity of earlier-hierarchy auditory regions, including belt cortex, produced significantly greater improvements in network integration and auditory-to-language routing efficiency than strengthening later-hierarchy regions. These findings provide novel measures for further investigation of auditory system dysfunction in schizophrenia, along with belt cortex as a mechanistically-grounded candidate for neuromodulatory intervention.

Several limitations should be noted. The predictions were not pre-registered; confirmation in larger independent cohorts with prospectively specified hypotheses will further establish the parcel-level mapping reported here. Most patients were receiving antipsychotic medication, which may influence resting-state FC; while convergence across multiple analytic approaches reduces this likelihood, a medication-naive sample would provide a cleaner test. As a cross-sectional resting-state study, the findings are correlational, and perturbation analyses speak to network sensitivity rather than causal mechanism [51, 56]. Finally, the study examined FC between predefined auditory and language parcels and did not characterize other networks implicated in AVH, such as the default mode, salience, and executive control networks.

## Acknowledgements

This work was supported by NIMH 2T32MH018870-38 and the Leon Levy Foundation to ACM, and R01MH49334 to DCJ and R01MH137769 to DCJ and AM. ACM and DCJ designed the research; ACM performed the research; ACM, AM, MA, YC, BY, JS, and GHP collected or prepared the data; ACM, YC, BY, AM, GHP, and DCJ prepared the manuscript. An LLM was used for copyediting.

