## Supplement for "Multiscale Network Disruption of Auditory-Language Circuits Underlying Auditory Hallucinations in Schizophrenia"

### Supplementary Materials
Multiscale Auditory–Language Disruption Underlying Hallucinations in Schizophrenia

Andrew C Murphy MD PhD[1,2,*], Antigona Martinez PhD[1,3], Yadi Chen MA[1], Michael Avissar MD PhD[4], Bahar Hazal Yalçınkaya PhD[2], Juan Sanchez-Pena MS[2], Gaurav H Patel MD PhD[1,2], and Daniel C Javitt MD PhD[1,2,3]

[1]Department of Psychiatry, Columbia University, New York, NY, USA
[2]New York State Psychiatric Institute, New York, NY, USA
[3]Nathan Kline Institute for Psychiatric Research, Orangeburg, NY, USA
[4]Alto Neuroscience
[*]

#### 1 Supplementary Methods

Full details of MRI acquisition, preprocessing, and post-processing are provided below. These procedures are referenced in the main text (Methods section).

##### 1.1 MRI Pre-processing

Results included in this manuscript come from preprocessing performed using *fMRIPrep* 24.1.1 ([15]; [14]; RRID:SCR_016216), which is based on *Nipype* 1.8.6 ([24]; [26]; RRID:SCR_002502).

###### 1.1.1 Preprocessing of B0 inhomogeneity mappings

A total of 1 fieldmaps were found available within the input BIDS structure for this particular subject. A *B0* nonuniformity map (or *fieldmap*) was estimated from the phase-drift map(s) measure with two consecutive GRE (gradient-recalled echo) acquisitions. The corresponding phase-map(s) were phase-unwrapped with `prelude` (FSL None).

###### 1.1.2 Anatomical data preprocessing

A total of 1 T1-weighted (T1w) images were found within the input BIDS dataset. The T1w image was corrected for intensity non-uniformity (INU) with `N4BiasFieldCorrection` [54], distributed with ANTs 2.5.3 [2, RRID:SCR_004757], and used as T1w-reference throughout the workflow. The T1w-reference was then skull-stripped with a *Nipype* implementation of the `antsBrainExtraction.sh` workflow (from ANTs), using OASIS30ANTs as target template. Brain tissue segmentation of cerebrospinal fluid (CSF), white-matter (WM) and gray-matter (GM) was performed on the brain-extracted T1w using `fast` [FSL (version unknown), RRID:SCR_002823, 59]. Brain surfaces were reconstructed using `recon-all` [FreeSurfer 7.3.2, RRID:SCR_001847, 13], and the brain mask estimated previously was refined with a custom variation of the method to reconcile ANTs-derived and FreeSurfer-derived segmentations of the cortical gray-matter of Mindboggle [RRID:SCR_002438, 35]. A T2-weighted image was used to improve pial surface refinement. Volume-based spatial normalization to two standard spaces (MNI152NLin2009cAsym, MNI152NLin6Asym) was performed through nonlinear registration with `antsRegistration` (ANTs 2.5.3), using brain-extracted versions of both T1w reference and the T1w template. The following templates were selected for spatial normalization and accessed with *Ciric2022* [24.2.0, 7]: *ICBM 152 Nonlinear Asymmetrical template version 2009c* [[19], RRID:SCR_008796; TemplateFlow ID: MNI152NLin2009cAsym], *FSL's MNI ICBM 152 non-linear 6th Generation Asymmetric Average Brain Stereotaxic Registration Model* [[18], RRID:SCR_002823;

TemplateFlow ID: MNI152NLin6Asym]. *Grayordinate* "dscalar" files containing 91k samples were resampled onto fsLR using the Connectome Workbench [20].

##### 1.1.3 Functional data preprocessing

For each of the 1 BOLD runs found per subject (across all tasks and sessions), the following preprocessing was performed. First, a reference volume was generated, using a custom methodology of *fMRIPrep*, for use in head motion correction. Head-motion parameters with respect to the BOLD reference (transformation matrices, and six corresponding rotation and translation parameters) are estimated before any spatiotemporal filtering using `Jenkinson2002` [FSL, 31]. The estimated *fieldmap* was then aligned with rigid-registration to the target EPI (echo-planar imaging) reference run. The BOLD reference was then co-registered to the T1w reference using `bbregister` (FreeSurfer) which implements boundary-based registration [27]. Co-registration was configured with six degrees of freedom. Several confounding time-series were calculated based on the *preprocessed BOLD*: framewise displacement (FD), DVARS and three region-wise global signals. FD was computed using two formulations following Power (absolute sum of relative motions, [43]) and Jenkinson (relative root mean square displacement between affines, [31]). Additionally, a set of physiological regressors were extracted to allow for component-based noise correction [*CompCor*, 4]. The confound time series derived from head motion estimates and global signals were expanded with the inclusion of temporal derivatives and quadratic terms for each [47]. Frames that exceeded a threshold of 0.5 mm FD or 1.5 standardized DVARS were annotated as motion outliers. Additional nuisance timeseries are calculated by means of principal components analysis of the signal found within a thin band (*crown*) of voxels around the edge of the brain, as proposed by [41]. The BOLD time-series were resampled onto the left/right-symmetric template "fsLR" using the Connectome Workbench [20].

#### 1.2 MRI Post-processing

The eXtensible Connectivity Pipeline-DCAN (XCP-D) [38, 9, 48] was used to post-process the outputs of *fMRIPrep* version 24.1.1 [16, 17, RRID:SCR_016216]. XCP-D was built with *Nipype* version 1.9.2 [25, RRID:SCR_002502].

**Segmentations** The following atlases were used in the workflow: the Schaefer Supplemented with Subcortical Structures (4S) atlas [50, 42, 34, 40, 21] at 10 different resolutions, the Glasser atlas [22], the Gordon atlas [23], the Tian subcortical atlas [52], the HCP CIFTI subcortical atlas [21], the MIDB precision brain atlas [29], and the Myers-Labonte infant atlas [39].

**Confound regression** Framewise displacement was calculated from the motion parameters using the formula from Power et al. [44], with a head radius of 50 mm. In total, 24 nuisance regressors were selected from the preprocessing confounds, according to the '24P' strategy [8, 48]. The BOLD data were converted to NIfTI format, despiked with *AFNI*'s *3dDespike*, and converted back to CIFTI format. Nuisance regressors were regressed from the BOLD data using a denoising method based on *Nilearn*'s approach. The timeseries were band-pass filtered using a second-order Butterworth filter, retaining signals between 0.01–0.08 Hz. The denoised BOLD was smoothed using *Connectome Workbench* with a Gaussian kernel (FWHM=6 mm).

The amplitude of low-frequency fluctuation (ALFF) [60] was computed using the Lomb-Scargle periodogram [36, 49, 53, 51]. Regional homogeneity (ReHo) [33] was computed using surface-based *2dReHo* [58]. Processed functional timeseries were extracted from residual BOLD using Connectome Workbench [37].

Many internal operations of *XCP-D* use *AFNI* [11, 12], *Connectome Workbench* [37], *ANTS* [3], *TemplateFlow* version 24.2.2 [10], *matplotlib* version 3.10.0 [30], *Nibabel* version 5.3.2 [6], *Nilearn* version 0.11.1 [1], *numpy* version 2.2.1 [28], *pybids* version 0.18.1 [57], and *scipy* version 1.15.1 [56].

#### 1.3 Network construction

Functional brain networks were constructed from denoised BOLD time series in surface (grayordinate) space. Cortical time series were parcellated using the Glasser multimodal cortical atlas, which defines 360 cortical parcels based on multimodal structural and functional features. For each parcel, a representative regional time series was obtained by averaging the BOLD signal across all vertices assigned to that parcel.

Functional connectivity between parcels was quantified using Pearson correlation coefficients computed between pairs of parcel-level time series, we did not apply thresholding due to the importance of weak connections in emergent network dynamics [46]. Correlation coefficients were Fisher $r$-to-$z$ transformed prior to statistical analysis to improve normality. Subject-level functional connectivity matrices were constructed using these Fisher-transformed correlation values and treated as weighted, undirected networks.

Subsequent analyses focused on connections between parcels belonging to the Cole–Anticevic auditory and language networks [32], with within-network connections excluded where specified to isolate cross-network interactions.

### 2 Statistical analyses

Associations between network-derived metrics and AVH severity were assessed using multinomial regression with PANSS P3 as the outcome variable. For each analysis, the regression coefficient ($\beta$) corresponding to the network metric of interest was taken as the estimated direction and magnitude of the association.

Statistical significance of multinomial regression coefficients was evaluated using nonparametric permutation testing (10,000 permutations), with two-sided $p$ values computed as the proportion of permuted coefficients with absolute value greater than or equal to the observed coefficient.

Associations between network metrics were assessed using Pearson correlation coefficients, with significance evaluated using permutation testing (10,000 permutations). Group comparisons were evaluated using permutation tests on raw group differences or difference-of-differences statistics. For pathway-specific analyses, permutation-based null models were constructed by randomly reassigning parcels to pathway roles while preserving pathway cardinality. *In silico* perturbation analyses were conducted by uniformly scaling all connections incident on a given parcel and recomputing network measures across a continuous spectrum spanning connection weakening (applied to HC data) through strengthening (applied to Scz data). For surface-based analyses, spatial autocorrelation was accounted for using spin-test null models [55]. Multiple comparisons were controlled using the false discovery rate (FDR).

### 3 Network metrics

Auditory network community structure was identified for each subject using modularity maximization (Louvain algorithm), which partitions the network into groups of densely interconnected regions. From this partition, we computed *assortativity*, a measure of how sharply segregated the communities are, with higher values indicating greater internal fragmentation of the auditory network. We computed *pathway efficiency* using mean first passage time from random-walk dynamics, capturing how efficiently signals travel from auditory source regions through belt cortex to language targets. For each parcel, we computed an *incident total-weight* measure summarizing the cumulative strength of its significant connections, and a *participation coefficient* quantifying the degree to which a region's connections span multiple communities, distinguishing hub regions that bridge communities from those confined within a single community. Full mathematical definitions and computational details are provided in the Supplementary Methods.

#### 3.1 Community detection in the auditory network

Community structure was identified separately for each subject using modularity maximization with the Louvain algorithm [5] ($\gamma = 1.0$), repeated 200 times per subject. The final partition was defined as the partition with the highest mean Rand $z$-score relative to all other partitions, selecting the solution maximally consistent with the ensemble.

##### 3.2 Auditory network assortativity

Assortativity was computed from the subject-specific community partition as:

$$\text{Assortativity} = \frac{\langle w_{\text{within}} \rangle - \langle w_{\text{between}} \rangle}{\langle w_{\text{within}} \rangle},$$

where $\langle w_{\text{within}} \rangle$ and $\langle w_{\text{between}} \rangle$ denote mean within- and between-community functional connectivity, respectively. Higher values indicate greater network segregation.

##### 3.3 Auditory–language pathway efficiency

Pathway efficiency was computed using mean first passage time (MFPT, [45]) from random-walk dynamics, restricted to the auditory–language subgraph. For each source–target pair, efficiency was defined as the inverse MFPT:

$$\text{Efficiency}_{i \to j} = \frac{1}{\text{MFPT}_{i \to j}}.$$

Pathway efficiency was summarized by averaging across specified auditory source, relay, and language target parcels with directionality preserved (auditory → language).

##### 3.4 Incident total-weight measures

The incident total weight for each parcel was defined as the sum of all significant (FDR- and spin-test-corrected) multinomial regression coefficients for edges incident on that parcel. Pairwise differences between parcel incident weights were assessed using permutation testing in which parcel labels were randomly shuffled to generate null distributions.

##### 3.5 Participation coefficient

The participation coefficient [45] of parcel $i$ was defined as:

$$PC_i = 1 - \sum_{m=1}^{M} \left( \frac{k_{i,m}}{k_i} \right)^2$$

where $k_i$ is the total weighted degree of parcel $i$ and $k_{i,m}$ is the sum of edge weights to parcels in community $m$. Values approaching 0 indicate provincial hubs; values approaching 1 indicate connector hubs. Associations between participation coefficients and auditory network assortativity were assessed using Pearson correlation with permutation-based significance testing, and pairwise differences between parcels were evaluated using permutation-based pairwise tests.

#### References

[1] Alexandre Abraham, Fabian Pedregosa, Michael Eickenberg, Philippe Gervais, Andreas Mueller, Jean Kossaifi, Alexandre Gramfort, Bertrand Thirion, and Gaël Varoquaux. Machine learning for neuroimaging with scikit-learn. *Frontiers in neuroinformatics*, page 14, 2014.

[2] B AVANTS, C EPSTEIN, M GROSSMAN, and J GEE. Symmetric diffeomorphic image registration with cross-correlation: Evaluating automated labeling of elderly and neurodegenerative brain. *Medical Image Analysis*, 12:26–41, 2 2008. ISSN 13618415. doi: 10.1016/j.media.2007.06.004.

[3] Brian B Avants, Nick Tustison, Gang Song, et al. Advanced normalization tools (ants). *Insight j*, 2 (365):1–35, 2009.

[4] Yashar Behzadi, Khaled Restom, Joy Liau, and Thomas T Liu. A component based noise correction method (compcor) for bold and perfusion based fmri. *NeuroImage*, 37:90–101, 2007. ISSN 1053-8119. doi: 10.1016/j.neuroimage.2007.04.042. URL https://www.sciencedirect.com/science/article/pii/S1053811907003837.

[5] Richard F Betzel. Community detection in network neuroscience. pages 149–171, 2023.

[6] Matthew Brett, Christopher J. Markiewicz, Michael Hanke, Marc-Alexandre Côté, Ben Cipollini, Paul McCarthy, Dorota Jarecka, Christopher P. Cheng, Yaroslav O. Halchenko, Michiel Cottaar, Eric Larson, Satrajit Ghosh, Demian Wassermann, Stephan Gerhard, Gregory R. Lee, Hao-Ting Wang, Erik Kastman, Jakub Kaczmarzyk, Roberto Guidotti, Jonathan Daniel, Or Duek, Ariel Rokem, Cindee Madison, Brendan Moloney, Félix C. Morency, Mathias Goncalves, Ross Markello, Cameron Riddell, Anibal Sólon, Christopher Burns, Jarrod Millman, Alexandre Gramfort, Jaakko Leppäkangas, Jasper J.F. van den Bosch, Robert D. Vincent, Henry Braun, Krish Subramaniam, Dimitri Papadopoulos Orfanos, Andrew Van, Krzysztof J. Gorgolewski, Pradeep Reddy Raamana, Julian Klug, B. Nolan Nichols, Eric M. Baker, Soichi Hayashi, Basile Pinsard, Christian Haselgrove, Mark Hymers, Oscar Esteban, Serge Koudoro, Fernando Pérez-García, Jérôme Dockès, Nikolaas N. Oosterhof, Bago Amirbekian, Ian Nimmo-Smith, Ly Nguyen, Samir Reddigari, Samuel St-Jean, Egor Panfilov, Eleftherios Garyfallidis, Gael Varoquaux, Jon Haitz Legarreta, Kevin S. Hahn, Lea Waller, Oliver P. Hinds, Bennet Fauber, Jacob Roberts, Jean-Baptiste Poline, Jon Stutters, Kesshi Jordan, Matthew Cieslak, Miguel Estevan Moreno, Tomáš Hrnčiar, Valentin Haenel, Yannick Schwartz, Zvi Baratz, Benjamin C Darwin, Bertrand Thirion, Carl Gauthier, Igor Solovey, Ivan Gonzalez, Jath Palasubramaniam, Justin Lecher, Katrin Leinweber, Konstantinos Raktivan, Markéta Calábková, Peter Fischer, Philippe Gervais, Syam Gadde, Thomas Ballinger, Thomas Roos, Venkateswara Reddy Reddam, and freec84. nipy/nibabel:, June 2022. URL `https://doi.org/10.5281/zenodo.591597`.

[26] Krzysztof J Gorgolewski, Oscar Esteban, Christopher J Markiewicz, Erik Ziegler, David Gage Ellis, Michael Philipp Notter, Dorota Jarecka, Hans Johnson, Christopher Burns, Alexandre Manhães-Savio,

Carlo Hamalainen, Benjamin Yvernault, Taylor Salo, Kesshi Jordan, Mathias Goncalves, Michael Waskom, Daniel Clark, Jason Wong, Fred Loney, Marc Modat, Blake E Dewey, Cindee Madison, Matteo di Oleggio Castello, Michael G Clark, Michael Dayan, Dav Clark, Anisha Keshavan, Basile Pinsard, Alexandre Gramfort, Shoshana Berleant, Dylan M Nielson, Salma Bougacha, Gael Varoquaux, Ben Cipollini, Ross Markello, Ariel Rokem, Brendan Moloney, Yaroslav O Halchenko, Wassermann Demian, Michael Hanke, Christian Horea, Jakub Kaczmarzyk, Gilles de Hollander, Elizabeth DuPre, Ashley Gillman, David Mordom, Colin Buchanan, Rosalia Tungaraza, Wolfgang M Pauli, Shariq Iqbal, Sharad Sikka, Matteo Mancini, Yannick Schwartz, Ian B Malone, Mathieu Dubois, Caroline Frohlich, David Welch, Jessica Forbes, James Kent, Aimi Watanabe, Chad Cumba, Julia M Huntenburg, Erik Kastman, B Nolan Nichols, Arman Eshaghi, Daniel Ginsburg, Alexander Schaefer, Benjamin Acland, Steven Giavasis, Jens Kleesiek, Drew Erickson, René Küttner, Christian Haselgrove, Carlos Correa, Ali Ghayoor, Franz Liem, Jarrod Millman, Daniel Haehn, Jeff Lai, Dale Zhou, Ross Blair, Tristan Glatard, Mandy Renfro, Siqi Liu, Ari E Kahn, Fernando Pérez-García, William Triplett, Leonie Lampe, Jörg Stadler, Xiang-Zhen Kong, Michael Hallquist, Andrey Chetverikov, John Salvatore, Anne Park, Russell Poldrack, R Cameron Craddock, Souheil Inati, Oliver Hinds, Gavin Cooper, L Nathan Perkins, Ana Marina, Aaron Mattfeld, Maxime Noel, Lukas Snoek, K Matsubara, Brian Cheung, Simon Rothmei, Sebastian Urchs, Joke Durnez, Fred Mertz, Daniel Geisler, Andrew Floren, Stephan Gerhard, Paul Sharp, Miguel Molina-Romero, Alejandro Weinstein, William Broderick, Victor Saase, Sami Kristian Andberg, Robbert Harms, Kai Schlamp, Jaime Arias, Dimitri Papadopoulos Orfanos, Claire Tarbert, Arielle Tambini, Alejandro De La Vega, Thomas Nickson, Matthew Brett, Marcel Falkiewicz, Kornelius Podranski, Janosch Linkersdörfer, Guillaume Flandin, Eduard Ort, Dmitry Shachnev, Daniel McNamee, Andrew Davison, Jan Varada, Isaac Schwabacher, John Pellman, Martin Perez-Guevara, Ranjeet Khanuja, Nicolas Pannetier, Conor McDermottroe, and Satrajit Ghosh. Nipype. *Software*, 2018. doi: 10.5281/zenodo.596855.

[27] Douglas N Greve and Bruce Fischl. Accurate and robust brain image alignment using boundary-based registration. *NeuroImage*, 48:63–72, 2009. ISSN 1095-9572. doi: 10.1016/j.neuroimage.2009.06.060.

[28] Charles R. Harris, Jarrod K. Millman, Stéfan J. van der Walt, Ralf Gommers, Pauli Virtanen, David Cournapeau, Eric Wieser, Julian Taylor, Sebastian Berg, Nathaniel J. Smith, Robert Kern, Matti Picus, Stephan Hoyer, Marten H. van Kerkwijk, Matthew Brett, Allan Haldane, Jaime Fernández del Río, Mark Wiebe, Pearu Peterson, Pierre Gérard-Marchant, Kevin Sheppard, Tyler Reddy, Warren Weckesser, Hameer Abbasi, Christoph Gohlke, and Travis E. Oliphant. Array programming with NumPy. *Nature*, 585(7825):357–362, September 2020. ISSN 1476-4687. doi: 10.1038/s41586-020-2649-2. URL `https://www.nature.com/articles/s41586-020-2649-2`. Bandiera_abtest: a computer-science;software;solar-physics.

[51] Paul A Taylor, Gang Chen, Daniel R Glen, Richard C Reynolds, and Robert W Cox. Lomb-scargle your way to rsfc parameter estimation in afni-fatcat. *International Society for Magnetic Resonance in Medicine*, 2018. Description of Lomb-Scargle applied to scrubbed fMRI data in AFNI.

[52] Ye Tian, Daniel S Margulies, Michael Breakspear, and Andrew Zalesky. Topographic organization of the human subcortex unveiled with functional connectivity gradients. *Nature neuroscience*, 23(11):1421–1432, 2020. doi: 10.1038/s41593-020-00711-6. URL `https://doi.org/10.1038/s41593-020-00711-6`.

[53] RHD Townsend. Fast calculation of the lomb–scargle periodogram using graphics processing units. *The Astrophysical Journal Supplement Series*, 191(2):247, 2010.

[54] N J Tustison, B B Avants, P A Cook, Y Zheng, A Egan, P A Yushkevich, and J C Gee. N4itk: Improved n3 bias correction. *IEEE Transactions on Medical Imaging*, 29:1310–1320, 2010. ISSN 0278-0062. doi: 10.1109/TMI.2010.2046908.

[55] František Váša, Jakob Seidlitz, Rafael Romero-Garcia, Kirstie J. Whitaker, Gideon Rosenthal, Petra E. Vértes, Maxwell Shinn, Aaron Alexander-Bloch, Peter Fonagy, Raymond J. Dolan, Ian M. Goodyer, Olaf Sporns, and Edward T. Bullmore. Adolescent tuning of association cortex in human structural brain networks. *Cerebral Cortex*, 28(1):281–294, 2017. doi: 10.1093/cercor/bhx101.

[56] Pauli Virtanen, Ralf Gommers, Travis E. Oliphant, Matt Haberland, Tyler Reddy, David Cournapeau, Evgeni Burovski, Pearu Peterson, Warren Weckesser, Jonathan Bright, Stéfan J. van der Walt, Matthew Brett, Joshua Wilson, K. Jarrod Millman, Nikolay Mayorov, Andrew R. J. Nelson, Eric Jones, Robert Kern, Eric Larson, C. J. Carey, İlhan Polat, Yu Feng, Eric W. Moore, Jake VanderPlas, Denis Laxalde, Josef Perktold, Robert Cimrman, Ian Henriksen, E. A. Quintero, Charles R. Harris, Anne M. Archibald, Antônio H. Ribeiro, Fabian Pedregosa, and Paul van Mulbregt. SciPy 1.0: fundamental algorithms for scientific computing in Python. *Nature Methods*, 17(3):261–272, March 2020. ISSN 1548-7105. doi: 10.1038/s41592-019-0686-2. URL `https://www.nature.com/articles/s41592-019-0686-2`. Bandiera_abtest:.

[57] Tal Yarkoni, Christopher J Markiewicz, Alejandro de la Vega, Krzysztof J Gorgolewski, Taylor Salo, Yaroslav O Halchenko, Quinten McNamara, Krista DeStasio, Jean-Baptiste Poline, Dmitry Petrov, et al. Pybids: Python tools for bids datasets. *Journal of open source software*, 4(40), 2019.

[58] Bo Zhang, Fei Wang, Hao-Ming Dong, Xiao-Wei Jiang, Sheng-Nan Wei, Miao Chang, Zhi-Yang Yin, Ning Yang, Xi-Nian Zuo, Yan-Qing Tang, and Ke Xu. Surface-based regional homogeneity in bipolar disorder: A resting-state fMRI study. *Psychiatry Research*, 278:199–204, August 2019. ISSN 0165-1781. doi: 10.1016/j.psychres.2019.05.045. URL `http://www.sciencedirect.com/science/article/pii/S0165178119305384`.

[59] Y Zhang, M Brady, and S Smith. Segmentation of brain mr images through a hidden markov random field model and the expectation-maximization algorithm. *IEEE Transactions on Medical Imaging*, 20: 45–57, 2001. ISSN 0278-0062. doi: 10.1109/42.906424.

[60] Qi-Hong Zou, Chao-Zhe Zhu, Yihong Yang, Xi-Nian Zuo, Xiang-Yu Long, Qing-Jiu Cao, Yu-Feng Wang, and Yu-Feng Zang. An improved approach to detection of amplitude of low-frequency fluctuation (ALFF) for resting-state fMRI: fractional ALFF. *Journal of Neuroscience Methods*, 172(1):137–141, July 2008. ISSN 0165-0270. doi: 10.1016/j.jneumeth.2008.04.012.
